# The global distribution of human-visible flower color diversity

**DOI:** 10.64898/2026.10.26.762001

**Authors:** James Boyko, Miranda A. Sinnott-Armstrong, Agnes S. Dellinger, Michael Foisy, Marisa M. Laitinen, Elia Schramek, Marjorie Weber, Thais Vasconcelos

## Abstract

The spectacular diversity of flower colors is among the most widely appreciated features of the natural world. For over a century, biologists and naturalists have speculated about whether human-visible flower color diversity shifts predictably across ecological systems worldwide. Yet challenges in measuring color across the vast diversity of flowering plants have limited global tests of these patterns. Using a computer-vision enabled analysis of community science data across 64,000 species, we modeled the major global features of human-visible flower color. We find a robust latitudinal gradient in color diversity after standardizing the number of species sampled from each flora, with flower colors occurring in more even proportions toward the equator. Modeling flower color alongside climate and pollination strategy reveals associations with both biotic and abiotic factors. These findings provide a global basis for testing how ecological and evolutionary processes shape the relative representation of floral phenotypes, beyond differences in species richness.

## Main

Across landscapes and through time, flowers have evolved an extraordinary bouquet of colors. This variation shapes how plants interact with their pollinators and, in turn, influences their reproductive success and diversity (Fenster et al., 2004). Foundational naturalists as early as Humboldt, Darwin, and Wallace speculated about variation in the human-visible spectrum of flower color across biomes and continents (Humboldt and Bonpland, 1809; Darwin, 1839; Wallace, 1878). Yet, the vast scale of flowering plant diversity has precluded global-scale analysis of flower color diversity. While powerful case-studies point to patterns in individual regions or taxonomic groups (e.g., Shrestha et al., 2013, Dalrymple et al., 2015; 2020; Gray et al., 2018; Dellinger et al., 2026), the emergent global features of flower-color diversity remain unresolved.

Traits related to ecological interactions, such as flower color, have long been predicted to increase in diversity towards the tropics, where the relative impact of selection should be expected to shift away from abiotic environmental filtering (i.e., the harsh climate of the poles) to more biotically driven selection of the species-rich equatorial regions (Schemske, 2009). Approximately 90% of flowering plant species depend on animals for pollination (Ollerton et al., 2011; Stephens et al., 2023), and flower color– even when limited to the human-visible spectrum–is well-known to mediate interactions with key animal pollinators (Sprengel, 1793). Through this lens, flower color diversity may be expected to generally increase towards the tropics across continents and clades. However, while the relative strength of selection from biotic interactions may increase in general towards the tropics, the major pollinator groups–including bees, butterflies, and birds–occur in both tropical and temperate regions, and their relative abundance, species richness, and phenotypic diversity varies considerably across geography and subclades (e.g., Rahbek and Graves, 2000; Heinrich et al., 2026). These differences may impose distinct selective pressures on different flowering plant species, generating geographic variation in floral coloration within and across plants that obfuscates the emergence of a general latitudinal pattern.

Adding further complexity, growing evidence also points to abiotic factors as underappreciated drivers of selection on flower-color, with theoretical implications for global color-diversity predictions. Beyond producing visible colors, floral pigments can protect sensitive reproductive tissues from cold temperatures (Chalker-Scott, 2008; Steyn et al., 2009), ultraviolet radiation (Jordan, 2004; Gould, 2004), and water stress (Vaidya et al., 2018). Outside of flowers, latitudinal gradients in color darkness and lightness are known to be plentiful in nature (e.g., in fungi (Krah et al 2019); insects (Lopez et al 2021); birds (Delhey 2017, Delhey 2019); flowers (Koski and Ashman 2015)), and are often explained by ecogeographic rules such as Gloger’s Rule (darker where warm and wet, Gloger, 1833; Delhey, 2017; Delhey, 2019) or Bogert’s Rule (darker where cold, Bogert, 1949). These physiological functions of plant pigments may favor the evolution of particular floral colors under particular environmental conditions and, in turn, shape the emergent composition and diversity of local flower color on a landscape. Recent studies focused on particular regions or clades have produced mixed results, with abiotic stress promoting flower color diversity in some (Dellinger et al., 2026), and constraining that diversity through environmental filtering in others (Gray et al., 2015).

Ultimately, whether the complex interplay of abiotic and biotic selection on floral coloration in species and local communities scales up to a detectable emergent pattern of flower-color diversity at the global scale remains unclear. Limited taxonomic and geographic coverage of flower-color data has historically hindered efforts (e.g., Dalrymple et al., 2015; Dellinger et al., 2025), as compiling standardized data for flower color at a large spatial and phylogenetic scale poses substantial challenges. Spectrophotometric measurements are costly and time-consuming to collect across tens of thousands of species, while data from herbarium specimen labels can be hard to standardize broadly since the original flower color has faded. Photographs vary in lighting conditions and require visible colors to be assigned consistently to discrete categories (Bergeron and Fuller, 2018; Valenta et al., 2021). However, modern advances in computer vision technology (e.g. McKenzie et al., 2025) combined with the rapid growth of global community-science photograph data have now laid the groundwork for the development of methods to characterize the main flower color of hundreds of thousands of observations spanning tens of thousands of species at a global-scale.

Here, we develop a computer vision pipeline to score standardized flower color data from photographs and apply it to community-science resources, assembling a global flower-color dataset comprising more than 64,000 species (Supplementary Methods S1; Table S1). Our standardized dataset represents roughly 20% of flowering plant diversity and more than five times the species coverage of the largest comparable dataset (∼11,000 species in McKenzie et al., 2025). We scored visible-spectrum flower colors from approximately 733,000 images from over 500,000 georeferenced, research-grade iNaturalist observations, spanning 64 plant orders, 396 families, 7,901 genera, and latitudes from 62°S to 83°N, assigning each species to one of ten color categories: white, yellow, purple, pink, cream, orange, red, blue, green and brown-maroon. We evaluated the image-derived color labels against independent, blinded human annotations (Supplementary Methods S2; Fig. S1; Table S2). We apply the resulting dataset to model geographic variation in flower color composition, test whether flower color diversity follows a robust latitudinal gradient, and examine how emergent global patterns relate to abiotic conditions and plant pollination strategies.

### A global evaluation of human-visible flower-color composition

Examining the data together provides a generalized look at the composition and variation of flower colors worldwide. White and yellow dominated globally, together accounting for 59% of species (33% and 26%, respectively), while blue, green and brown-maroon were the rarest color categories, each below 2% (Fig. 1D). While every color category occurred on every continent except Antarctica, their prevalence shifted notably with geography. Red and orange each made up about 0.5% of species in Europe but 4.4% and 5.2% in South America, whereas blue was most common in Europe and temperate Asia (Fig. S2). Color composition also differed sharply among families. White dominated Rosaceae, Myrtaceae, Rubiaceae and Caryophyllaceae (54–66%), while yellow was the leading color in Asteraceae (48%) and Fabaceae (39%), and purple was most common in Lamiaceae (41%) and Plantaginaceae (40%). Notably, some lineages and floras drew on a narrow palette. For example, more than 90% of species in Hypericaceae and Dilleniaceae were yellow, and two thirds of Caryophyllaceae were white.Interestingly, families known for their specialized pollination systems exhibited more diversified color profiles; Bromeliaceae and Orchidaceae, for instance, had species spread across all ten categories, with no color exceeding 28%. Among regions with at least 50 species assigned a color category, the South Island of New Zealand had the lowest rarefied effective color diversity (the exponential of mean Shannon entropy) with 61% of species white-flowered. Several small tropical island floras, including the Galápagos, were also among the most lowest rarefied effective color diversity, while the most color-diverse floras were found in Ecuador, Peru, Colombia, Peninsular Malaysia, Borneo and tropical Africa (see full dataset in Table S3).

**Fig 1.**
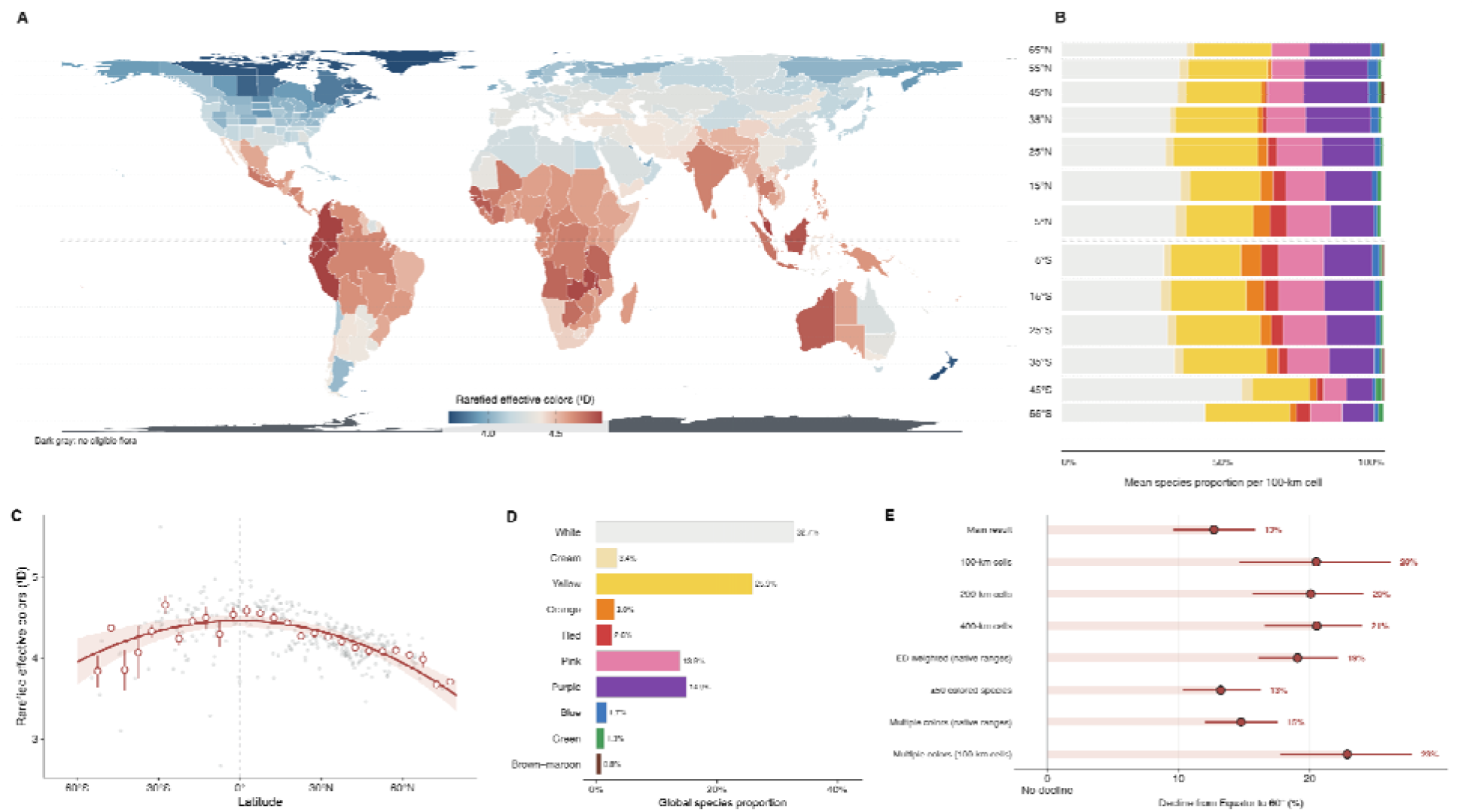
Human-visible flower-color diversity declines poleward across native regional floras. (A) Flower-color diversity across 351 native regional floras, standardized to samples of 20 species. Dark gray indicates regions without eligible floras (Supplementary Methods S3). (B) Mean proportions of species in each color category across 100-km occurrence cells, grouped into 10° latitude bands and aligned with the map (Supplementary Methods S3). (C) Regional flower-color diversity against latitude, with 5°-bin means ± SE. The curve is a descriptive quadratic fit; shading shows its 95% spatial-bootstrap confidence band (Supplementary Methods S3). (D) Global proportions of species assigned to each color category (Supplementary Methods S1). (E) Estimated percentage decline in flower-color diversity from the equator to 60° absolute latitude under the main analysis and robustness checks, including evolutionary-distinctiveness weighting. Bars show 95% spatial-bootstrap intervals (Supplementary Methods S3-S4).

### Flower color diversity increased toward the equator

Mapping the relative frequencies of flower colors worldwide revealed a latitudinal gradient in flower color diversity, paralleling latitudinal patterns in species richness (Jablonski et al., 2006; Brown, 2013) and color diversity in other organisms and plant structures (Wallace, 1878; Bailey, 1978; Adams et al., 2014; Sinnott-Armstrong et al., 2018; Cooney et al., 2022; Dellinger et al., 2026). Flower colors occurred in more even proportions near the equator, with rarified effective color diversity (exponential of mean Shannon entropy) 12.7% lower at 60° absolute latitude compared to 0° (95% spatial-bootstrap CI: 9.6–15.8%; Fig. 1A,C; Supplementary Methods S3; Table S4). This decline reflected increasing dominance of common flower colors, particularly white and purple, at higher latitudes, whereas tropical floras contained greater proportions of less common colors such as pink, red, and orange leading to an overall more evenly diverse composition (Fig. 1B; Table S4). A visual comparison across Whittaker climate biomes was consistent with the pattern, with tropical rainforests presenting the highest flower color diversity, followed by tropical seasonal forests and savannas (Supplementary Methods S3; Fig. S3; Table S5).

This latitudinal signal in flower color diversity was robust to common confounders of global diversity patterns. When floras were standardized to the same number of species, tropical assemblages still distributed those species more evenly among color categories (Fig. 1E; Table S4), indicating that the pattern is not simply a consequence of the tropics containing more flowering plant taxa. The gradient was also evident in occurrence-derived 100-km assemblages (Fig. S4; Table S4). Accounting for phylogenetic sampling biases by weighting species by their evolutionary distinctiveness similarly left the richness-corrected relationship with absolute latitude largely unchanged (Spearman’s ρ = −0.688, compared with −0.728 before phylogenetic weighting; Fig. S5; Table S4), suggesting that the gradient was not driven by the disproportionate sampling of species-rich clades sharing similar colors in certain regions. Finally, the latitudinal pattern also persisted after controlling for geographic variation in sampling, the exclusion of poorly observed species, and allowing species to retain multiple commonly observed colors rather than assigning each a single dominant color (Fig. 1E; Supplementary Methods S4; Fig. S5; Table S4). Together, these analyses confirm what some early naturalists suspected, that the floras of the tropics are indeed exceptional in their floral chromatic diversity, beyond what is expected simply from their species richness alone.

While the latitudinal gradient in flower color diversity provides an emergent geographic signature of the underlying ecological and evolutionary processes impacting color diversity on a global scale, latitude itself cannot be interpreted as a direct mechanism driving biological processes. Instead, multiple factors that can covary with latitude, including climatic conditions and pollinator communities (Sakhalkar et al. 2026), are predicted to drive the evolution and ecological filtering of floral traits, including investment into flower color and consequently flower color diversity. To explore signatures consistent with underlying biological drivers of geographic patterns, we explicitly modeled flower color alongside local climate data and plant pollination mode.

### Climatic predictors of flower-color categories

We first considered whether the distributions of individual flower colors were associated with species’ climatic niches. We modeled the occurrence of each of the ten flower colors across the 37,643 species shared between our dataset and the largest currently available phylogeny of flowering plants (Carruthers et al., 2026) as a function of six climatic variables known to affect the synthesis of colorful pigments in flowers (maximum temperature, actual evapotranspiration, vapor pressure deficit, solar radiation, wind speed and UV-B) in ten separate one color-versus-rest phylogenetic logistic regressions (Supplementary Methods S5; Fig. S6; Table S6). We found that accounting for the evolutionary relationships among species was particularly important for disentangling evolutionarily robust patterns from raw color-climate correlations. Estimates from phylogenetic and non-phylogenetic models were only moderately correlated (*r* = 0.63), and 11 of the 60 estimated relationships reversed direction when phylogenetic relationships were included (Fig. 2; Table S6), suggesting visually apparent color-climate associations often reflect shared evolutionary history rather than independent responses to climate.

**Fig 2.**
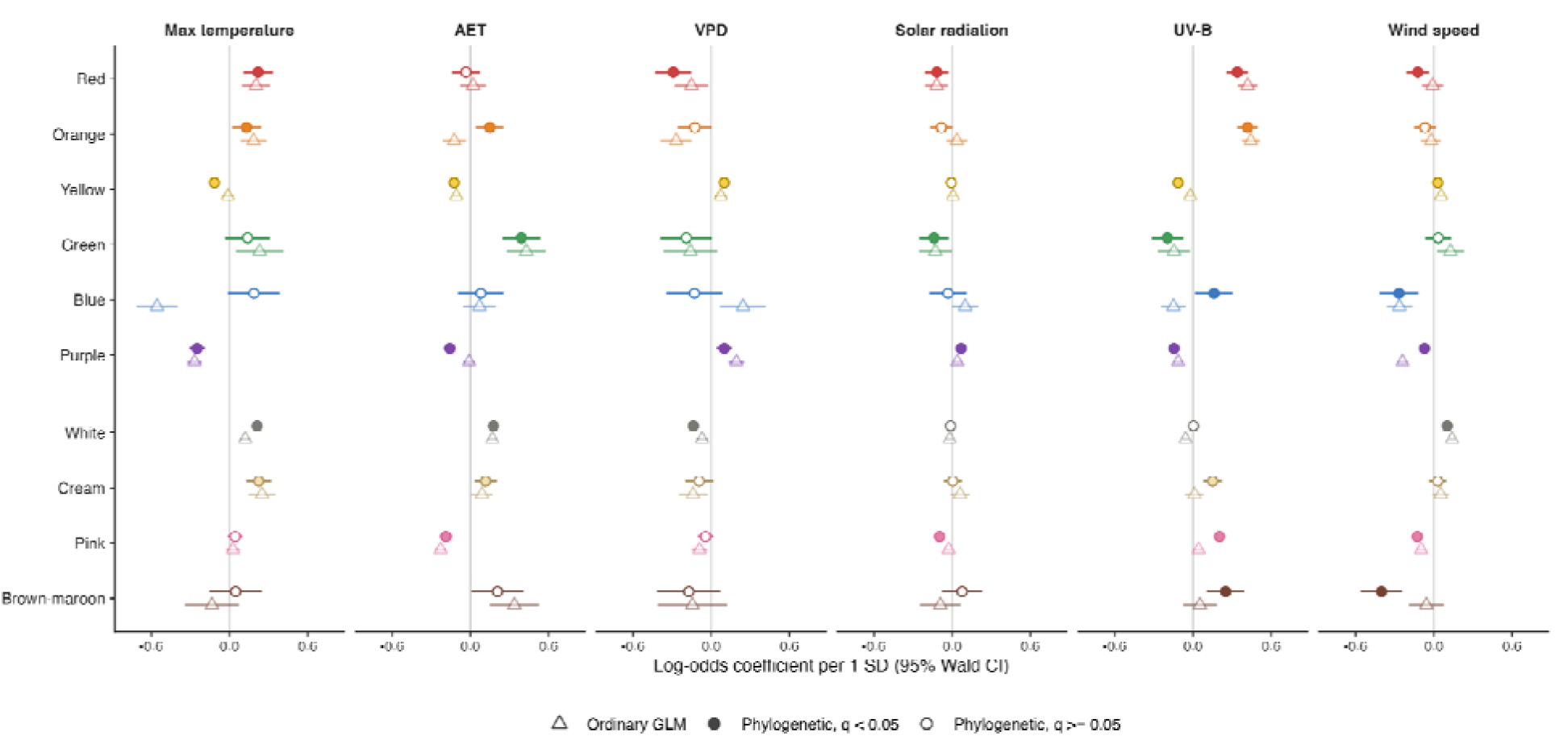
Species color–climate associations with and without phylogenetic dependence. All models use 37,643 species with complete values of six standardized climatic-niche positions and the same species color labels. Coefficients for ten one-versus-rest responses, with 95% Wald intervals. Open triangles show ordinary logistic estimates; circles show phylogenetic maximum penalized-likelihood estimates, filled for q < 0.05 and open otherwise. False-discovery correction covers all 60 non-intercept tests in the primary model family. Zero lines indicate no conditional association. Max temperature, monthly mean maximum temperature; AET, monthly total actual evapotranspiration; VPD, monthly mean vapor pressure deficit; Solar radiation, monthly mean downward surface shortwave radiation; UV-B, annual mean ultraviolet-B radiation from glUV; Wind speed, monthly mean wind speed. Species-level climatic-niche positions are medians across occupied 100-km cells (Supplementary Methods S5).

When accounting for phylogeny, we found individual flower colors were robustly associated with several distinct climatic conditions. Among the most common was UV-B - a measure of the ultraviolet-B radiation reaching the ground - which displayed phylogenetically-robust and significant associations with nine of the ten colors (Fig. 2; Table S6). Species in high-UV-B environments were more likely to have red (β = 0.34), orange (β = 0.42), brown-maroon (β = 0.25), cream (β = 0.15), pink (β = 0.20) or blue flowers (β = 0.16), but less likely to have green (β = −0.20), yellow (β = −0.11) or purple flowers (β = −0.15), suggesting that UV-driven processes may play an underappreciated role in shaping global patterns of color from the tropics to the poles where UV-B is lowest. Indeed, many of the colors that are more common in high-UV-B environments (e.g., red, orange) are often, but not exclusively, produced by carotenoids that can be protective of the photosynthetic apparatus (Young, 1991) which can be damaged by excessive UV-B radiation (Teramura and Sullivan, 1994). However, UV-B was strongly correlated with absolute latitude (r = −0.91). After including absolute latitude in the phylogenetic models, only the UV-B association with blue remained significant after false-discovery correction. Purple (produced almost exclusively by the common plant pigments anthocyanins and, in some Caryophyllales, betalains) was more common in cooler climates (β = -0.25 with maximum temperature), consistent with a reported protective benefit of anthocyanins against cold stress (Steyn et al., 2008), and yellow, which can often be produced by carotenoid pigments, was more prevalent in drier habitats (β = 0.10 with vapor pressure deficit), perhaps reflecting the dual protective role of carotenoids against against UV-B and water stress. These large-scale correlational relationships link the climatic niches of species to their flower colors, and point to testable mechanistic predictions about how the physiological consequences of certain colors and their underlying pigments translate to large-scale geographic patterns.

Because different climatic variables were associated with the spatial distributions of individual colors, we tested whether climate alone could reproduce the observed latitudinal gradient in flower color diversity. We modeled the color composition of each grid cell as a function of its climate, without latitude as a predictor, and averaged predictions over the months in which species in that cell were recorded flowering (Supplementary Methods S6; Fig. S7). Projecting the fitted model onto all grid cells for which climate data were available yielded a global surface of expected flower-color composition (Fig. 3A; 7,973 cells; Fig. S8; Table S7). Together, these globally-projected climatic associations reproduced the observed decline in flower color diversity toward the poles, predicting 19% fewer effective flower-color diversity at 60° than at the equator compared to an observed decline of 22% (Fig. 3C; Table S7). Changes in the frequencies of individual colors contributed unevenly to this decline. White and yellow, which together dominated assemblages across latitudes, changed little between 0–10° and >50° (from 0.33 to 0.37 and from 0.23 to 0.24, respectively). By contrast, the predicted frequencies of red and orange declined approximately fivefold (from 0.047 to 0.008 and from 0.049 to 0.010, respectively), while pink also declined and purple and blue increased (Fig. S9; Table S7). The predicted poleward decline in effective flower-color diversity primarily reflected decreasing frequencies of red and orange. Increasing frequencies of blue and purple partly offset this decline, while white contributed little to the overall trend (Table S7).. Together, this analysis reveals that global climate-color associations contain sufficient geographic structure to generate the broad global pattern without the need to explicitly incorporate latitude. Cell-level agreement between observed and climate-predicted effective color diversity was nonetheless modest (Pearson’s r = 0.46 across 2,653 fitting cells), and residuals showed geographic structure (Fig. 3B), indicating variation not captured by the climate model.

**Fig 3.**
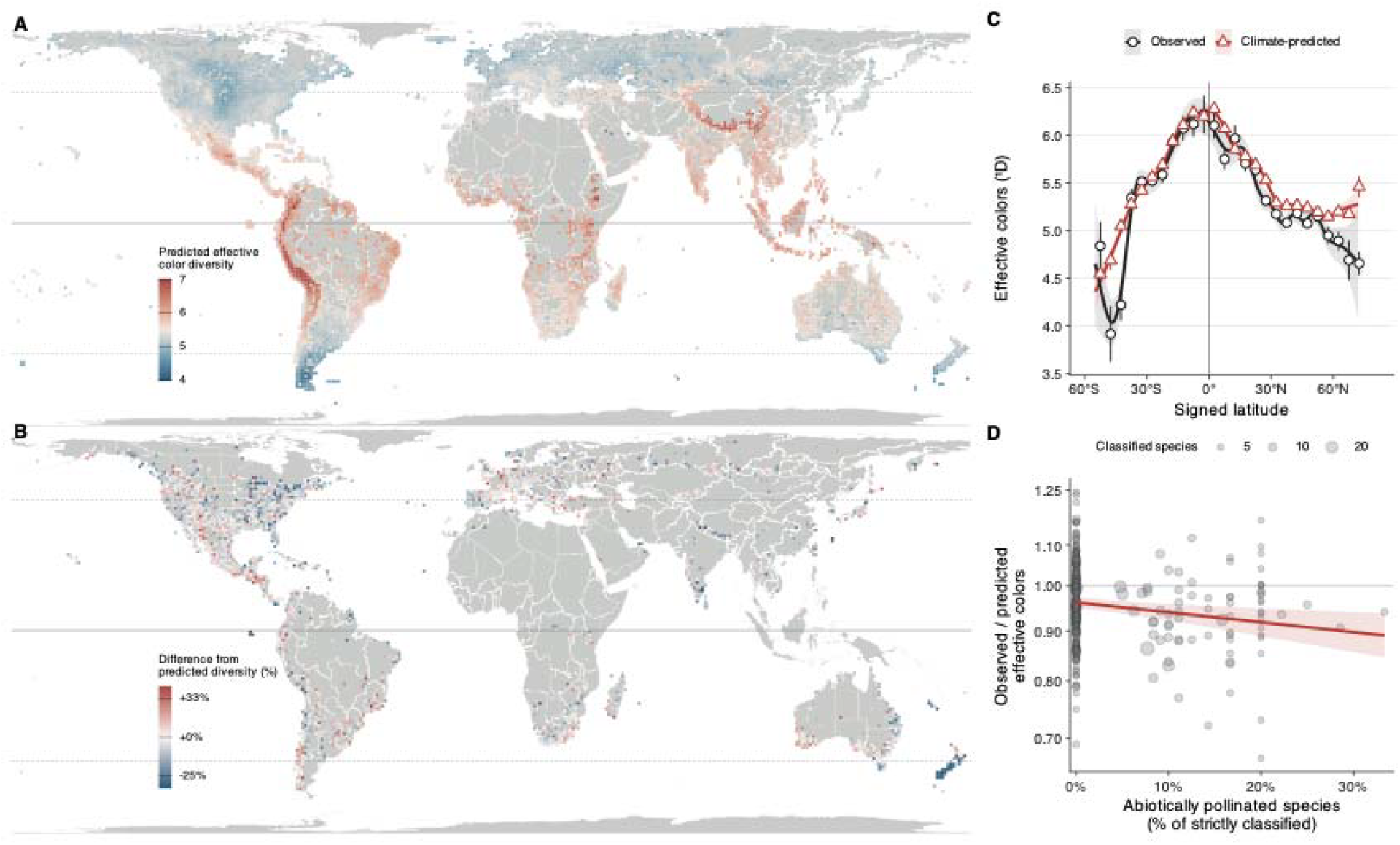
Climate-only predictions reproduce the broad latitudinal decline but leave residuals associated with abiotic pollination. (A) Predicted effective colors in 7,973 climate-bearing 100-km cells. A Dirichlet-multinomial model uses six climate variables without latitude; monthly predicted compositions are averaged with species-balanced flowering-month weights, then entropy is exponentiated (Supplementary Methods S6; Fig. S8). (B) Percentage difference between observed and climate-predicted effective color diversity across 2,653 fitting cells, calculated as 100 × (observed/predicted − 1). Colors follow a logarithmic ratio scale; positive values indicate higher diversity than predicted (Supplementary Methods S6; Table S7). (C) Observed and predicted effective color diversity against signed latitude, with 5°-bin means ± SE and descriptive GAM smooths with 95% intervals. Separate regressions against absolute latitude estimate 22% observed and 19% predicted declines from the equator to 60° (Table S7). (D) Latitude-adjusted observed/predicted ratio against the abiotically pollinated percentage of species classified by Stephens et al. (2023), in 315 cells with ≥5 classified species. Point area shows classified species. The weighted fit, with 1,000-km block-clustered 95% interval, gives 2.3% fewer effective colors per 10 percentage points (95% CI 0.8–3.8%; sensitivity: Fig. S10). Classified species represent a median 3.6% of colored species per cell.

### Pollination mode helps explain flower-color diversity beyond climate

Despite the overall agreement between climate predictions and observed flower-color diversity, several conspicuous mismatches (Fig. 3B) suggest that other factors, including the abundance and diversity of animal pollinators, also contribute to the observed pattern. The diversity of animal pollinators and animal-pollinated plants (Ollerton et al., 2026) as well as disparity in other (non-color) floral traits (Chartier et al., 2021), varies considerably across latitude and are generally thought to increase in tropical and subtropical zones, where we also observe the most even representation of flower colors. Ideally, global analyses of flower color diversity would model climate alongside plant pollination mode and the local diversity and abundance of the relevant pollinator community to explore their relative importance in structuring the distribution of flower colors. Unfortunately, the complexities of global pollinator diversity, encompassing many distinct lineages of insects as well as vertebrates spanning specialists to generalists (Sakhalkar et al., 2026), means that global-scale maps of plant-specific pollinator diversity do not exist, precluding a direct test comparable to our above climatic analysis. However, data on pollination mode, and specifically whether plants are mainly animal- or abiotically-pollinated, is more readily available (Stephens et al., 2023). To evaluate whether a signature of animal-pollination mode on flower-color diversity emerges at the global scale, we cross-referenced our data with a recent dataset of pollination mode for 1,201 flowering plant taxa (Stephens et al., 2023), and modelled the residual variation from our climate model as a function of the fraction of abiotically pollinated species in each grid cell, adjusting for absolute latitude (Supplementary Methods S7; Table S8). This analysis tests whether deviations from climate-only predictions are partly explained by the proportion of animal-pollinated plants in a region. If accounting for pollination mode strengthens climate-color associations, this would suggest that interactions with animal pollinators partially underlie global patterns of flower-color diversity.

We found that grid cells with a higher proportion of abiotically pollinated species had fewer effective flower colors than predicted by climate, with a 2.3% decrease for every 10-percentage-point increase in abiotic pollination (95% CI 0.8–3.8%; 315 cells; Fig. 3D; Fig. S10; Table S8). Thus, within the subset of species for which pollination-mode data were available, assemblages containing a greater representation of abiotically pollinated species had lower flower-color diversity than climate alone predicted (Fig. 3D). This analysis also recovered the previously reported relative increase in abiotically pollinated plants at higher latitudes, where wind-pollinated species – the predominant group within this category – are generally more common (Ollerton et al., 2011) (Fig. S10; Table S8). Further, several areas classically recognized as depauperate in pollinator diversity had fewer effective colors than expected based on their climate alone (e.g., Galápagos - 36% fewer, and New Zealand - 27% fewer, Fig. 3B; Tables S7-8). Together, climate–color associations reproduced much of the broad latitudinal gradient, while pollination mode was associated with departures from climate-predicted effective color diversity.

### Conclusions and Future directions

Flower color is among the most widely celebrated biological phenotypes within modern biodiversity, yet challenges with measuring color at scale have limited analyses of flower color to individual clades and regional case studies. Here, we leverage modern advances in automated computer vision and community-science data to create a large, standardized dataset of human-visible flower color across roughly 20% of flowering plant species. Geographic analysis of the dataset reveals a robust latitudinal gradient in flower color diversity, with colors occurring in increasingly uneven proportions further from the equator. These patterns correlate strongly with climatic variables, indicating that abiotic conditions are likely an important driver of both individual flower colors and flower color diversity. Improved predictive power when incorporating pollination-mode into models of climate-color-diversity associations further points to signatures consistent with both biotic and abiotic drivers shaping flower-color diversity across biomes.

Our results demonstrate the power of new computer vision-based methods to extract large trait datasets from citizen science observations. These new technologies allow biologists to document phenotypic diversity rapidly, consistently, and at unprecedented scales across the tree of life, revealing previously undetected patterns and providing crucial data to advance our understanding of global biodiversity.

Integrating computer-vision pipelines with more in-depth but scale-limited approaches will allow for even more powerful tests of specific hypotheses. For example, although more scale-limited, standardized field sampling across gradients can counter potential overrepresentation of human-conspicuous flowers in community science datasets, and spectrophotometer measurements can add information on color brightness, saturation, and reflectance in non-visible regions of the spectrum which cannot be measured from photographs alone. However, ultimately the computer vision-enabled big data approaches utilized here provide a powerful, standardized global picture of human-visible flower-color categories, providing a foundational synthesis of the emergent features of flower color diversity globally.

## Materials and Methods

### Flower color data

We obtained research-grade iNaturalist observations of vascular plants from GBIF (GBIF.org, 29 January 2026, https://doi.org/10.15468/dl.fgg926) and used images that PhenoVision (Dinnage et al. 2025) classified as showing flowers with high certainty, that carried a reusable Creative Commons license and that were identified at least to family. We selected up to 30 images per species with the highest PhenoVision probability, then up to 30 further observations per species from within its native range in the World Checklist of Vascular Plants (WCVP; Govaerts et al. 2021), drawn evenly across 10° × 10° latitude–longitude cells. Each image was submitted to GPT-5-nano (OpenAI) with a fixed prompt (text S1) in three runs. Some images supplied fewer than three usable answers and were recorded as incomplete responses (Table S1). The prompt asked for the predominant color of the flower petals or reproductive structures in one of ten human-visible categories (white, cream, yellow, orange, red, pink, purple, blue, green, brown–maroon) and whether the image was an open-flower close-up. We did not score ultraviolet reflectance, brightness or saturation, which cannot be standardized across community-science photographs.

We kept observations whose images were mostly judged open-flowered close-ups, because agreement with human annotators was higher in such images (Fig. S1; Table S2). We also kept only angiosperm observations located within the species’ native range, defined as the WGSRPD level-3 regions (Brummitt 2001) in which WCVP lists the species as native. The final dataset held 507,963 observations (733,278 images) of 64,263 species in 396 families (Table S1). Each species received a single dominant color, the category with the most votes summed over all of its runs, images and observations. One species received no color, leaving 64,262 species assigned to a color category, and all color percentages count species, not observations (Supplementary Methods S1).

### Color validation

We validated color scoring against independent human annotations of 1,000 images, sampled before curation with 100 per model-assigned dominant color. Each image was scored by two annotators who saw only the photograph and genus and could select alternative colors when uncertain. We compared agreement between annotators with agreement with the model’s image dominant color and observation color set (categories receiving more than 20% of its color votes). Agreement required overlapping color labels; images either annotator could not score were excluded from paired comparisons. We weighted rates by observing dominant-color frequencies in the final dataset and obtained 95% confidence intervals and paired differences from 10,000 bootstrap resamples within sampled color categories. The full protocol, unweighted and single-color comparisons, per-color performance, confusion patterns, image-filter and latitude checks, and an earlier unblinded round are in Supplementary Methods S2.

### Assemblage color diversity and latitude

Our primary analysis used native regional floras from the World Checklist of Vascular Plants (WCVP) in WGSRPD level-3 regions (Govaerts et al. 2021; Brummitt 2001). Observation names were resolved to accepted WCVP species, their color votes pooled, and each accepted species assigned one color. A species entered every region in which WCVP listed it as native. The analysis included 351 floras with at least 20 species, drawn from 63,847 accepted species with assigned flower colors. We measured diversity as the effective number of colors, ^1^D = exp(H), where H is Shannon entropy of species proportions across ten categories. We standardized richness by drawing 20 species without replacement 500 times per flora and exponentiating the mean entropy.

We fitted ln(rarefied ^1^D) = a + b|latitude| using polygon-centroid latitude and summarized the predicted decline from the equator to 60° as 100[1 − exp(60b)]. We obtained 95% percentile intervals from 999 bootstrap resamples of occupied 20° × 20° spatial blocks. A signed-latitude quadratic described the shape in Fig. 1C. A separate descriptive biome comparison pooled unique species within WorldClim– Whittaker biomes and calculated unrarefied effective colors (Fig. S3; Table S5). Assemblage construction, latitude definitions, metric distinctions, fitting and uncertainty are detailed in Supplementary Methods S3 and Table S4; sensitivity comparisons are defined in S4 (Fig. 1E; fig. S5; table S4).

### Robustness of the latitudinal gradient

We tested whether the gradient depended on sampling or on how species colors were assigned. We repeated the analysis with 100-, 200- and 400-km occurrence grids, adjusted for species richness and the fraction of each region’s native flora with color data, and raised the minimum flora size from 20 to 50 species with color assignments. We also rebuilt floras after excluding species with fewer than five or ten observations. Under an alternative color rule, each species’ contribution was split equally among colors receiving more than 20% of its votes.

To assess sensitivity to shared evolutionary history, we weighted species by evolutionary distinctiveness, each species’ share of the phylogeny’s branch lengths, and recalculated regional color diversity. We correlated richness-adjusted diversity with absolute latitude and compared it with the unweighted regional result. Only species present on the tree entered the weighted analysis. We also examined medians within latitude bands and compared equatorward and poleward floras within continents. Full definitions, samples, estimates and uncertainty are reported in Supplementary Methods S4.

### Species-level color–climate associations

We summarized each species’ climatic niche using the median climate of its uniquely occupied 100-km cells, giving each cell equal weight. Cell values were means of TerraClimate monthly values extracted at the locations, years and months of flowering observations (2006–2024), with annual mean glUV UV-B sampled at cell centers (Abatzoglou et al. 2018; Beckmann et al. 2014). The six predictors were maximum temperature, actual evapotranspiration, vapor pressure deficit, downward shortwave solar radiation, wind speed and UV-B.

We matched species to a recent phylogeny of flowering plants (Carruthers et al. 2026), retaining 37,643 tree-matched species with complete niche positions. For each discrete color category, we fitted a separate one-versus-rest phylogenetic logistic regression on all six predictors, standardized across these species to mean zero and standard deviation one. We used maximum penalized likelihood (phylolm::phyloglm, logistic_MPLE), optimizing the unchanged penalized likelihood with Subplex after diagnosing numerical instability (S5). We reported per-SD log-odds effects with 95% Wald intervals, and adjusted the 60 non-intercept tests by Benjamini–Hochberg false-discovery correction (q < 0.05).

Identical ordinary logistic models supplied the comparison in Fig. 2. Climate extraction, tree matching and repair, coverage, standardization, diagnostics and existing bound sensitivity are detailed in Supplementary Methods S5. Grid-level color composition is modeled separately.

### Grid-level climate model of color composition

To ask whether climate alone predicts the latitudinal gradient, we modeled the color composition of 100-km occurrence cells rather than individual species. The response of each cell was its number of species in each of the ten discrete color categories; each species counted once per cell. We used the 2,653 cells with at least 20 species with color assignments and cell climate values. Predictors were the six variables of the species models (maximum temperature, actual evapotranspiration, vapor pressure deficit, downward shortwave radiation, wind speed and UV-B), here as cell means of the observation-level values and standardized across the 2,653 cells. The model contained no latitude or other geographic term.

We fitted a Dirichlet-multinomial regression by maximum likelihood (MGLM 0.2.1; Kim et al. 2018), in which each color’s Dirichlet parameter is log-linear in the predictors (70 coefficients). We evaluated it with ten spatially contiguous blocks, refitting without each block and predicting it; one refit did not converge, leaving held-out predictions for 2,377 cells.

We applied the fitted model to the climate of each month in which flowering was recorded in a cell and averaged the predicted compositions over those months (Fig. 3). Each month was weighted by its number of species recorded in flower. We projected these predictions to all 7,973 cells with climate data. Diversity of the averaged composition was expressed as effective colors, exp(H), and compared with observed Miller–Madow-corrected diversity. Predicted and observed declines were estimated by regressing ln(^1^D) on absolute latitude over the same 2,653 cells and expressed as the percentage change from 0° to 60°. Residuals are log ratios of observed to predicted effective colors. The model, simulation check, cross-validation, residual analyses, extrapolation screen, color contributions and weighting alternatives are detailed in Supplementary Methods S6.

### Pollination mode and climate residuals

We took plant pollination modes from Stephens et al. (2023), who scored 1,201 angiosperm taxa as animal-, wind- or water-pollinated or a combination of these. We matched these taxa to our species by exact binomial after removing underscores, collapsing spaces and ignoring case, without resolving synonyms. Of 715 matched species, we classified 624 as biotically pollinated (animal only) and 47 as abiotically pollinated (wind, water or both). We excluded 40 species pollinated by both animals and wind or water, and 4 without a score.

The unit of analysis was the 100-km fitting cell of the grid-level climate model. For each cell we counted the strictly classified species recorded there and computed the fraction that were abiotically pollinated. The response was the cell’s Fig. 3B residual, the natural logarithm of the ratio of observed to climate-predicted effective colors. We analyzed the 315 cells with at least five strictly classified species. In these cells, classified species were a median of 3.6% of species, so the abiotic fraction describes the classified subset, not the full flora.

We regressed the residual on the abiotic fraction and absolute latitude by weighted least squares, with weights equal to the number of classified species. Standard errors were clustered by 1,000-km equal-area blocks (77 blocks), and 95% intervals are ±1.96 standard errors. A coefficient β on the abiotic fraction corresponds to a 100[exp(0.1β) − 1]% change in the observed-to-predicted ratio for each 10-percentage-point increase in the abiotic fraction. We repeated the analysis with minimum cell counts of three and ten classified species, and with classifications restricted to observed pollination systems or to confidence scores of at least 3 (Supplementary Methods S7).

